# SMC complexes can traverse physical roadblocks bigger than their ring size

**DOI:** 10.1101/2021.07.15.452501

**Authors:** Biswajit Pradhan, Roman Barth, Eugene Kim, Iain F. Davidson, Benedikt Bauer, Theo van Laar, Wayne Yang, Je-Kyung Ryu, Jaco van der Torre, Jan-Michael Peters, Cees Dekker

## Abstract

The ring-shaped structural-maintenance-of-chromosomes (SMC) complexes condensin and cohesin extrude loops of DNA as a key motif in chromosome organization. It remains, however, unclear how these SMC motor proteins can extrude DNA loops in chromatin that is bound with proteins. Here, using *in vitro* single-molecule visualization, we show that nucleosomes, RNA polymerase, and dCas9 pose virtually no barrier to DNA loop extrusion by yeast condensin. Strikingly, we find that even DNA-bound nanoparticles as large as 200 nm, much bigger than the SMC ring size, can be translocated into DNA loops during condensin-driven extrusion. Similarly, human cohesin can pass 200 nm particles during loop extrusion, which even occurs for a single-chain version of cohesin in which the ring-forming subunits are covalently linked and cannot open up to entrap DNA. These findings disqualify all common loop-extrusion models where DNA passes through the SMC rings (pseudo)topologically, and instead point to a nontopological mechanism for DNA loop extrusion.

**One-sentence summary:** Huge DNA-bound roadblocks can be incorporated into SMC-extruded DNA loops, pointing to a nontopological mechanism for loop extrusion.

---

Chromosome organization in eukaryotic cells is vital for genome segregation, gene regulation, and recombination^1,2^. Structural maintenance of chromosome (SMC) complexes such as condensin and cohesin play a key role in the threedimensional organization of the chromosome^3–5^. These SMC complexes are DNA-binding ATPases, in which a SMC heterodimer and kleisin subunit form a ring-like structure.^6,7^ SMC complexes organize the genome by extruding DNA loops in a processive and ATP-dependent manner^8–11^. Both cohesin and condensin have been shown to extrude loops *in vitro* on bare DNA molecules that were tethered onto a surface^12–16^.

It remains unclear, however, how SMC complexes deal with the abundant proteins that are part of chromatin in cells. *In vivo*, SMC complexes will encounter lots of DNA-binding proteins such as nucleosomes as well as large obstacles such as the DNA replisome, with a size of ∼20 nm^17^, or a transcribing RNA polymerase that, with its RNA transcript and spliceosome, can have a globular diameter of >70 nm^18^, i.e., even bigger than ∼35 nm ring size of SMCs. Some data indicate that SMCs can bypass DNA-bound objects, e.g. Kim *et al*.^14^ and Kong *et al*.^19^ found that nucleosomal DNA was compacted by cohesin and condensin, and yeast condensin was shown to be able to bypass another condensin to form higher-order DNA loops.^20^ Yet, other evidence has indicated a slowing down or blocking of DNA loop extrusion by specific proteins. For example, the DNA-binding protein CCCTC-binding factor (CTCF) regulates topologically associating domains and long-range interactions by blocking loop extrusion through specific binding to cohesin.^21^ Furthermore, *in vivo* data suggested that transcribing RNA polymerases are able to ‘push’ cohesin to the 3’-end of highly transcribed genes in eukaryotes^22,23^ and Brandão *et al*. theoretically predicted that transcription slows SMC complexes down but does not block SMC translocation in *B. subtilis*^24^. Under conditions in which cohesin diffuses along DNA, SMCs were reported to be blocked by objects larger than 20 nm.^25,26^ Overall, it therefore remains unresolved whether loop-extruding SMC complexes will block, stall, bypass, dissociate, or push obstacles that are present on the DNA upon encounter.

One might expect that this relates to the topology of the SMC complex during DNA loop extrusion. SMCs can embrace DNA in their ring structure, as cohesin was found to mediate sister chromatid cohesion through topological entrapment^27^ while condensin was reported to topologically link chromatid arms for their structural rigidity in mitosis^28^. Whether topological entrapment is necessary for loop extrusion is under debate. While so far all modelling^29–36^ assumes some form of DNA entrapment, recent data on human cohesin with a covalently closed-ring structure indicated its ability to extrude loops^13^, suggesting a possible pseudo- or non-topological loading of the SMC complex during DNA loop extrusion, where, respectively, the DNA gets inserted into the SMC ring without opening the ring, or where the DNA loop does not get embraced by the SMC ring at all but DNA binding occurs externally.

Here, we systematically study the effect of DNA-binding proteins on DNA-loop extrusion for both yeast condensin and (covalently closed) human cohesin. Strikingly, we find that particles as large as 200 nm, i.e., much bigger than the ∼35 nm condensin ring size, can be translocated into DNA loops during extrusion. This demonstrates that SMCs can handle protein-loaded chromatin substrates for loop extrusion without any problems, and provides direct evidence for a nontopological mechanism of SMC-driven DNA loop extrusion.

Loop extrusion was visualized *in vitro* by monitoring a Alexa647 fluorescently labelled ‘roadblock’ protein on fluorescently labelled DNA (sytox orange (SxO); Fig. 1a). Roadblock proteins were bound to 48.5kb λ-DNA, which was anchored onto a streptavidin-coated passivated glass surface at both its ends through biotin linkers (fig. S1) and imaged with homebuilt HILO sheet microscopy. Upon addition of condensin (0.5 nM) and ATP (5 mM), fluorescence spots locally formed on DNA, whose intensity grew until the slack was removed between the two DNA ends. Inplane buffer flow perpendicular to the DNA confirmed that these were extruded DNA loops. Consistent with previous findings,^12^ condensin extruded a DNA loop asymmetrically, and consequently, the DNA-bound roadblock was reeled towards the loop or remained fixed with respect to the loop position (Fig. 1c, fig. S2). Events where the roadblock was moving towards the loop to subsequently co-localize with the loop were identified as an encounter and considered for further analysis.

**Fig. 1.**
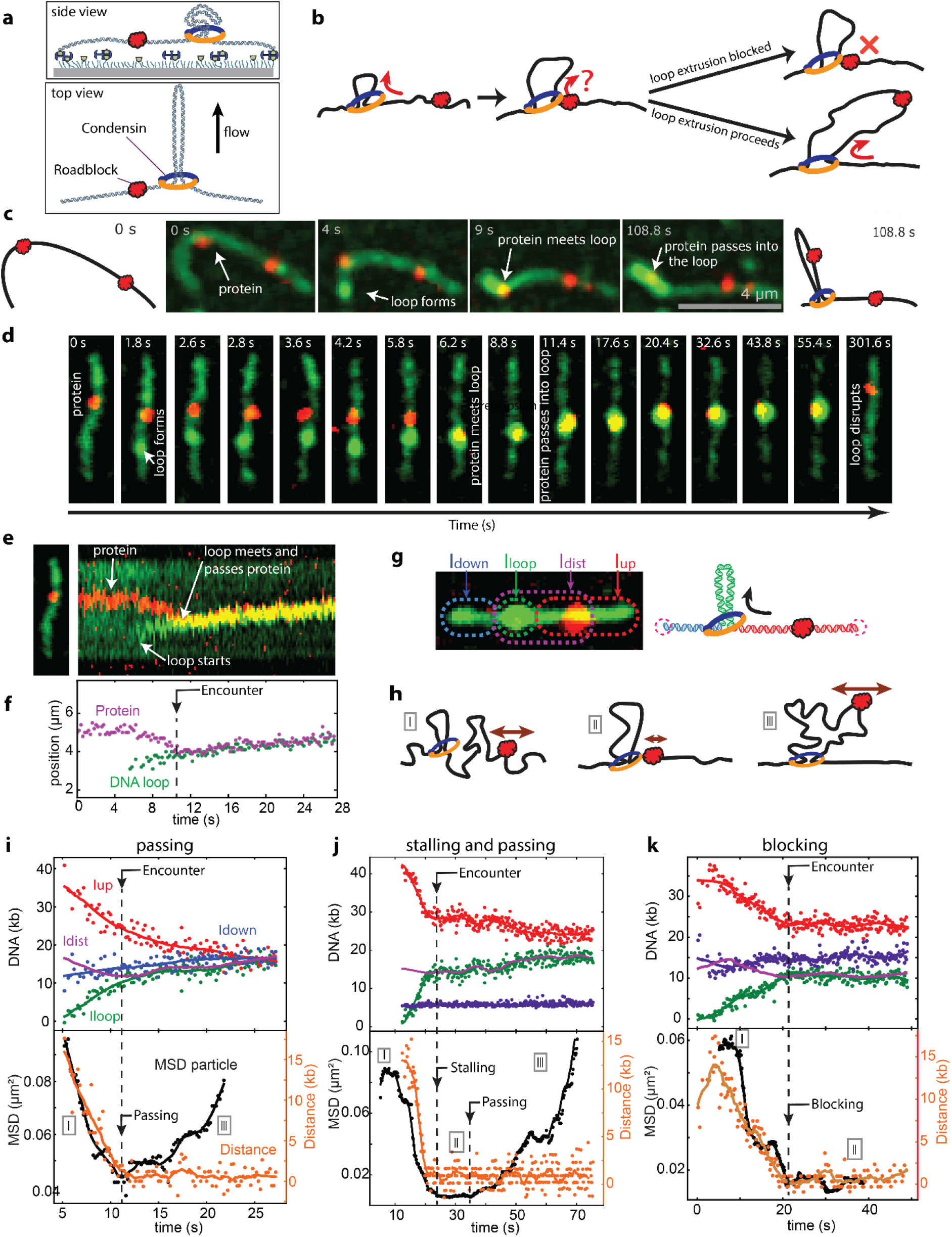
Visualization of a condensin encountering an obstacle while extruding a DNA loop. (a) Loop extrusion assay. (b) Two scenarios for condensin encountering an obstacle during loop extrusion: The obstacle can either pass into the loop or get blocked. (c) Snapshots of condensin bypassing a DNA-bound protein (dCas9) and including it in the loop during DNA loop extrusion with side flow. (d) Snapshots of DNA loop extrusion where the loop encounter accommodates a protein. (e) Kymograph for the event in (d). (f) Localizations of the protein (magenta) and DNA puncta (green) from the kymograph. (g) Color scheme denoting the various fluorescence intensities of different regions on DNA. (h) Cartoon showing the Brownian fluctuations of the roadblock at different stages of the encounter. (i) Kinetics of the DNA loop size formation for a passing event. Intensities are denoted in colors outlined in panel g. Bottom curve show MSD (moving average; black) and SMC-roadblock distance (orange). (j) Same as i, but for a stalling and subsequent passing event. (k) Same as i, but for a blocking event.

Upon encounter, loop extrusion by condensin either blocked, or the roadblock traversed the stem of the condensin-mediated DNA loop to subsequently translocate into the extruded DNA loop (Fig. 1b). Figure 1c shows an example of such a traversal, where the DNA formed a loop that grew in size until it encountered the roadblock (here dCas9) whereupon it continued to grow such that the loop encompassed the roadblock protein. When the loop spontaneously disrupted after 9 minutes, the protein was still bound to the DNA and, as expected, at its initial position. Alternatively, a roadblock can get blocked at the stem of the loop upon encounter (fig. S1). We never observed pushing of the roadblock along the DNA or disruption of the loop upon the encounter. To avoid imposing additional tension within the DNA by the side flow, which can slow down or stall loop extrusion^12,37^, most roadblock experiments were performed without side flow (Fig. 1d). Kymographs were used to visualize loop formation and roadblock positions over time, see Fig. 1e,f. Upon encounter, a continued movement of the now co-localized loop and roadblock signaled the passing of the roadblock into the loop, since the SMC still drove the loop expansion. Quantification of the loop size from the intensities (Fig. 1g) provided further details, see Fig. 1i,j,k for examples of a passage, stalling and passing, and blocking event, respectively.

We found that the mean square displacement (MSD) of the roadblock along the DNA serves as a good marker for discriminating whether the roadblock got blocked or passed into the extruded loop. For a passage event, the MSD (Fig. 1i, bottom) could be seen to rapidly decrease until the condensin-roadblock encounter, and rapidly increasing again after the encounter. This is explained from, respectively, the tightening up of the DNA between its two ends in the early phase of loop extrusion (i.e., going from I to II in Fig.1h), and the increased distance from the roadblock (inside the loop) to the loop stem which is associated with an increased Brownian motion (III in Fig. 1h) since the DNA within the loop is not under tension. The MSD (Fig. 1i, bottom) changes much more distinctly over time than the loop size (Fig. 1i, top), making it a discriminating measure for inferring whether the roadblock did or did not get incorporated into the extruded DNA loop.

In probing the effects of roadblocks on condensin-driven DNA loop extrusion, we first examined nucleosomes, the basic building block of chromatin with a diameter of ∼11 nm (Fig. 2a). Nucleosomes were reconstituted from yeast histone octamers with Alexa647 on H3A (Methods), and assembled as single nucleosomes on DNA (Fig.2b; fig. S6), yielding 2-5 isolated nucleosomes per λ-DNA in our optical assay. Both passing and blocking events were observed (Fig. 2c–g), but passing events were much more frequent (88±4%, Fig. 2i). A small fraction of the encounters showed stalling and passing with pausing time of ∼10 s (fig. S10). Notably, the blocking events almost always occurred at the end of a loop extrusion where the forces within the DNA were approximately similar to the stalling force for condensin (fig. S4a). Other proteins with similar diameters, specifically RNA polymerase (RNAP) from *E. Coli*, and dCas9 (Fig. 2h), showed similar passing fractions as the nucleosome, namely 92±5 % and 87±8 %, respectively. This indicates that ∼10 nm-sized DNA-binding proteins can be readily accommodated into the extruded loop.

**Fig. 2.**
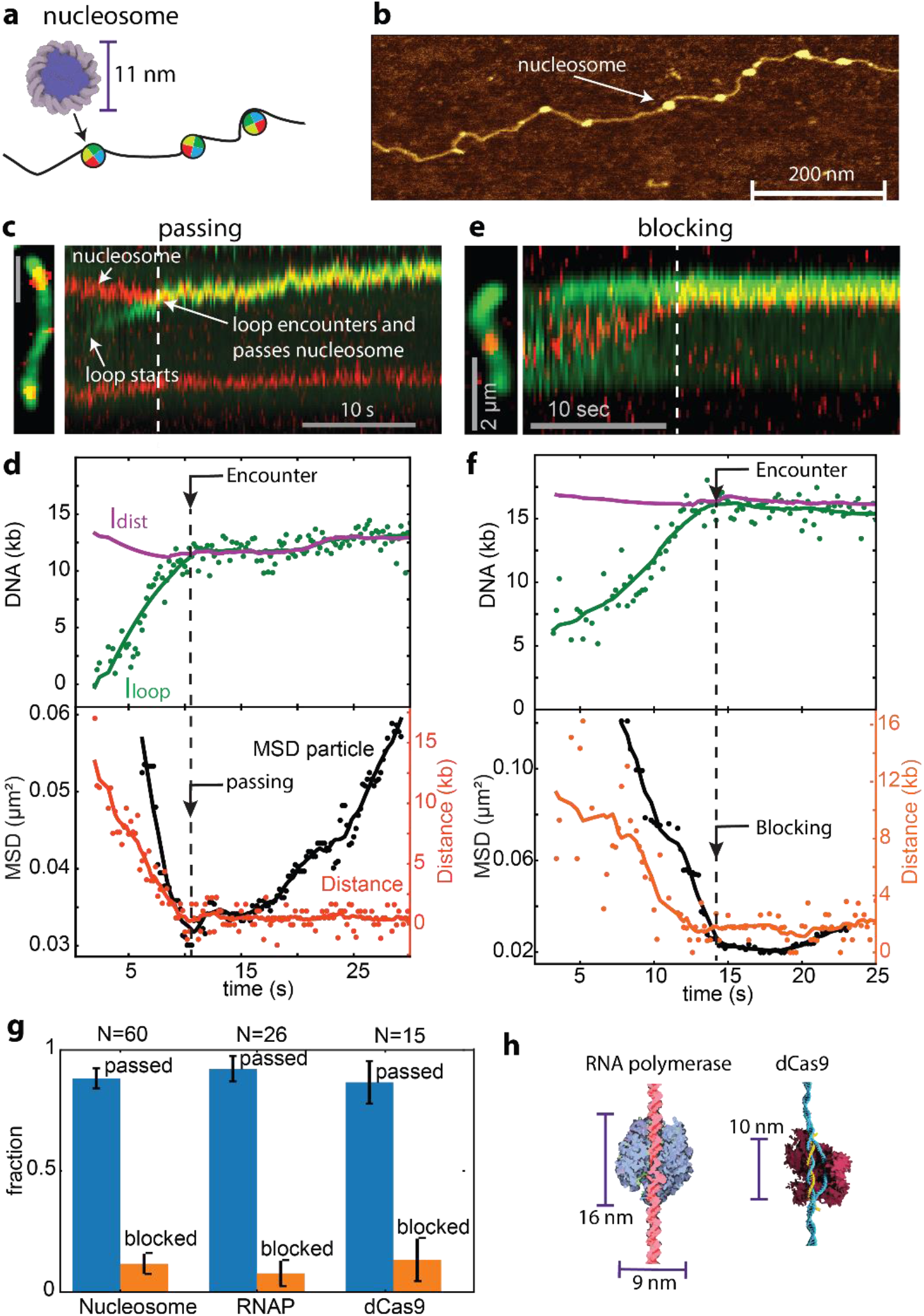
Loop-extruding condensin traverses nucleosomes and other DNA-bound proteins. (a) Schematic of nucleosomes on DNA. (b) AFM image of single nucleosomes on DNA. (c) Kymograph for a passage event. (d) Corresponding loop kinetics and MSD traces. (e, f) Same as c, d, but for a blocking event. (g) Fraction of the roadblocks passed (blue) and blocked (orange) for nucleosomes, RNAP and dCas9. (h) Schematics of RNAP and dCas9 (adapted from Wikimedia Commons).

Next, we asked if yet larger obstacles on DNA would block DNA-extruding condensin. Gold nanoparticles were chosen to controllably vary the size of the roadblocks, as they can be obtained with a narrow size distribution with a median diameter of 10–125 nm. Gold particles were functionalized with SH-PEG-NH2/SH-PEG-COOH (see Methods) which minimized interactions with DNA, condensin, and the glass surface. The amine (-NH2) groups facilitated binding to Alexa647 and to a dCas9-Snaptag that allowed sequence-specific anchoring of the nanoparticle to the DNA via dCas9. TEM imaging (Fig. 3b–d), which visualized the gold core but not the outer organic layer, showed a narrow size distribution. The hydrodynamic diameter was estimated from fluorescence correlation curves (fig. S5) and comparison of the TEM and hydrodynamic diameters yielded a thickness of 7.9±1.2 nm for the passivation layer. For the largest particle in our study, we used polystyrene nanoparticles with the same functionalization that added 16±2 nm to the 182±8 nm diameter as measured by EM, leading to a 198±10 nm total diameter of these ‘200 nm’ polystyrene nanoparticles.

**Fig. 3.**
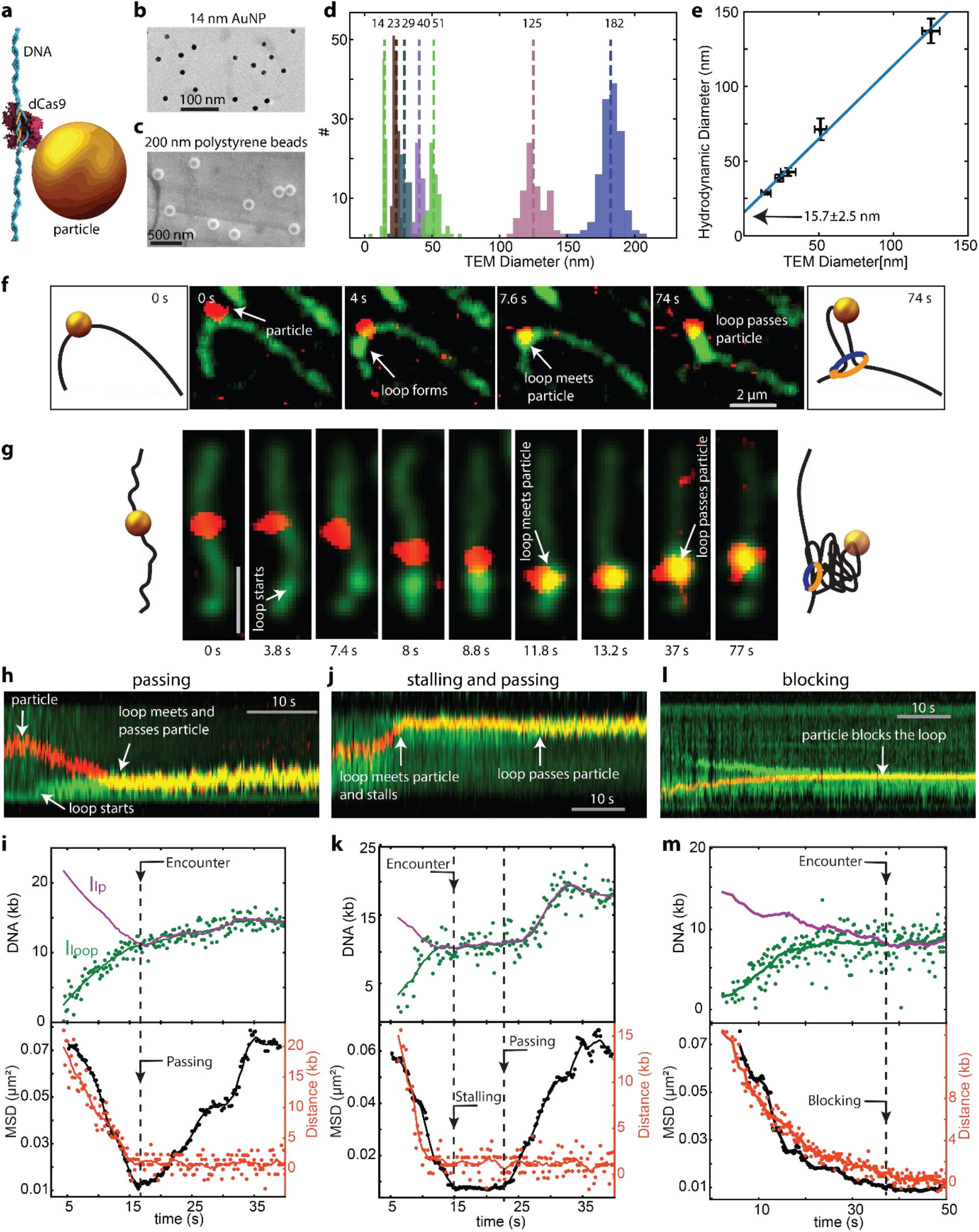
Functionalized gold nanoparticles as controlled steric obstacles. (a) Schematic showing the binding of a functionalized gold nanoparticle to the DNA through dCas9. (b) TEM image of functionalized gold nanoparticles of sizes 14 nm showing uniform sizes and absence of aggregates. (c) Idem for 186 nm polysterene beads. (d) Size distribution of the functionalized gold nanoparticles from the TEM images. (e) Correlation between hydrodynamic diameter obtained from FCS and diameters obtained from TEM. A linear fit shows an additional thickness of 15.7 nm on the nanoparticles. (f) Snapshots of DNA loop extrusion with side flow for a 39 nm gold nanoparticle (red) on the DNA (green) that ends up within the loop. (g) Snapshots of loop extrusion without side flow with a 39 nm gold nanoparticle (red) on the DNA (green). (h, j, l) Kymograph of DNA (green) and nanoparticle (red) corresponding to passing (h), stalling before passing (j) and blocking (l) events. (i, k, m) Loop kinetics and MSD traces corresponding to passing (i), stalling before passing (k) and blocking (m) events.

Condensin-driven loop extrusion was examined for functionalized nanoparticles of different sizes that were bound onto DNA. Figure 3f–m provides examples for gold nanoparticles with a hydrodynamic diameter of 39 nm. We observed different outcomes upon encounter of the condensin with the roadblock, viz., passing (Fig. 3f–i), stalling and passing (Fig. 3j,k, fig. S10), and blocking (Fig. 3l,m). Application of a side flow unequivocally demonstrated that a 39 nm particle can be inserted into the loop (Fig. 3f). From the statistics of passing versus blocking, 39 nm particles show similar high passing fractions (94±6%; including ‘stalling and passing’ events) as the nucleosomes (88±4%).

The most striking finding of our study is that DNA-bound particles with a diameter as large as 200 nm can pass into the loop during the process of DNA loop extrusion. Figure 4c shows representative side-flow images of DNA loop extrusion in the presence of a 200-nm particle where the particle can be seen moving towards the loop, then encountering the loop, to subsequently pass into the loop. Figs. 4c-e shows another example of a passage event where a 200 nm particle gets incorporated into the extruded DNA loop, as evidenced from the increase in both the loop size and the clear minimum in the MSD at the encounter. Notably, this 200 nm size is much larger than the condensin ring (cf. Fig. 4a for a comparison to scale), as the circular SMC complex has a diameter of about 35 nm^38^, as is e.g. evident from AFM imaging (fig. S6d, e), whereas theoretically one could imagine a ∼110 nm ring diameter (fig. S6f) when combining the ∼50 nm long smc arms and stretching the kleisin to its full maximal length, which however is nonphysiological.

**Fig. 4.**
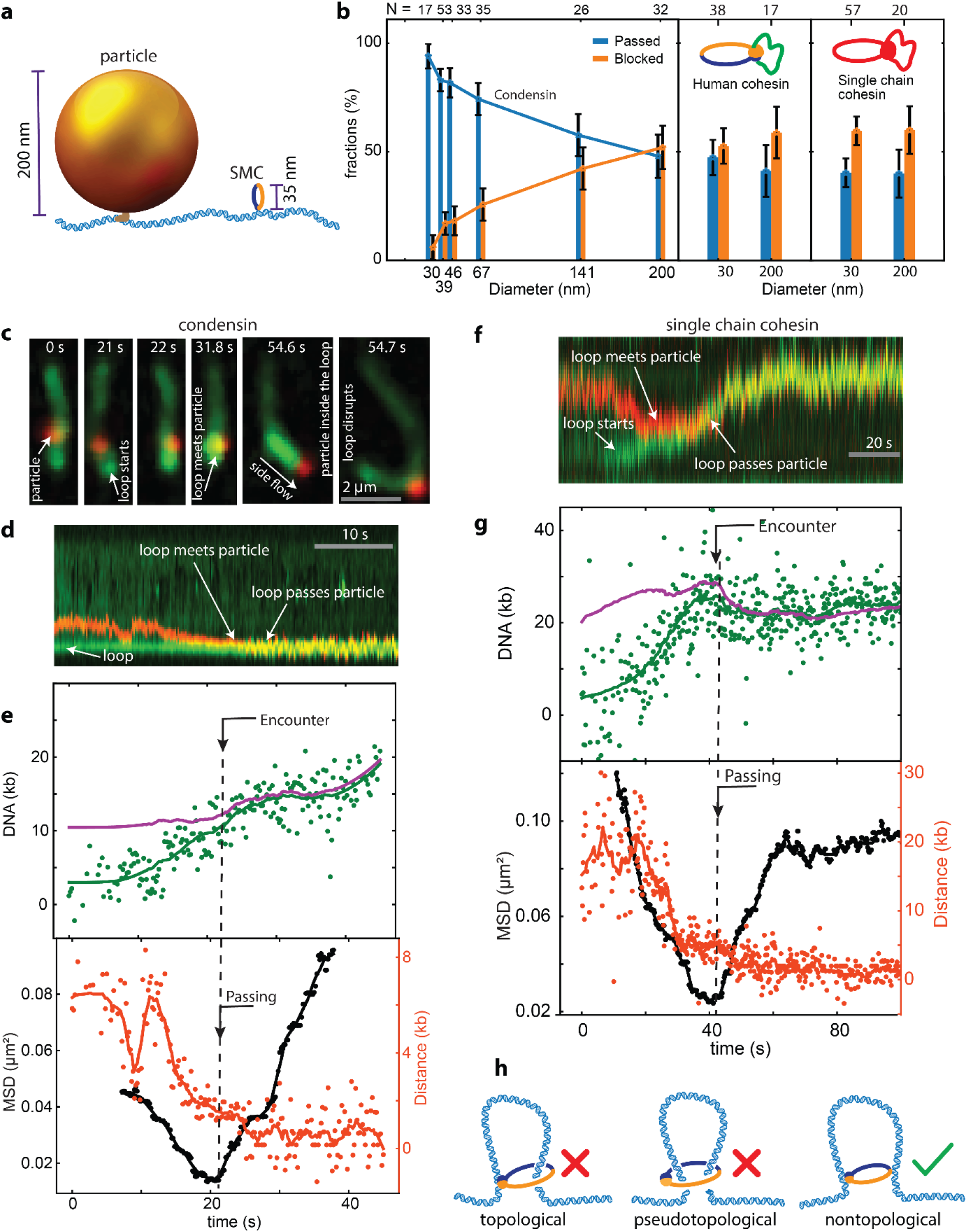
Condensin and cohesin can pass roadblocks bigger than their ring size. (a) Sketch of a Cas9-200nm nanoparticle and condensin on DNA, to scale. (b) Fraction of roadblocks passed into the loop or blocked by yeast condensin, human cohesin, and single-chain cohesin, for different particle sizes. (c) Snapshots of loop extrusion by condensin on a DNA that accommodates a 200 nm roadblock in a passing event. At 54.7 s, a side flow was applied and the particle can be seen inside the loop. (d) Kymograph of a passage event where condensin traverses a 200 nm roadblock. (e) Loop kinetics and MSD trace for the event in (d). (f) Kymograph of a passage event where single-chain human cohesin traverses a 200 nm roadblock. (g) Loop kinetics and MSD trace for event (f). (h) Models of topological, pseudotopological, and nontopological embracing of DNA underlying DNA loop extrusion.

As displayed in Fig.4b, the fraction of DNA-bound nanoparticles that traversed the condensin to get incorporated into the DNA loop decreased from >90 % for particle sizes up to 40 nm, to 53±8 % for the 200nm-size particles. The very large passing fractions for the smaller particles indicate that the vast majority of biologically relevant DNA-binding proteins will be readily accommodated into extruded DNA loops. While the passing fraction decreases with increasing particle size, it is remarkable that >50% of the 200 nm particles still get translocated into the extruded loop – implying that also very large protein complexes on DNA will traverse into DNA loops. About 10-20% of all encounters exhibited stalling before subsequent passing, with a characteristic stalling time of about 14 s (fig. S10).

To explore the generality of these findings, we performed similar experiments with human cohesin, which was also shown to extrude DNA loops.^13,14^ For both 30 nm and 200 nm DNA-bound nanoparticles, we observed that these roadblocks were able to pass the cohesin complex during loop extrusion. Figure 4g shows a typical example of a passage event that shows that cohesin behaves very similar to condensin, although the encounter/MSD analysis is slightly more involved due to the two-sided nature of loop extrusion for cohesin (see SI and fig. S7). Interestingly, both 30 nm and 200 nm particles traverse cohesin to pass into the loop with similar efficiencies, 47±8 % and 45±15 % respectively (Fig. 4b, right).

Our observation that 200-nm particles can be translocated into DNA loops by extruding SMC complexes suggests that the DNA that is being extruded is not encompassed by the ∼35 nm SMC ring structure. Alternatively, it is possible that large obstacles get translocated across an SMC complex by transiently opening the SMC ring structure. To rigorously test this important question, we examined a single-chain variant of human cohesin in the roadblock assay. In this engineered version of cohesin (fig. S6c), the three ring-forming subunits SMC3, SCC1 and SMC1 are expressed as a single fused polypeptide chain. Similar to wildtype cohesin, this fusion protein folds into a ring-like structure in which SMC1 and SMC3 hetero-dimerize via the hinge domain to close the ring. This hinge-hinge interaction is furthermore crosslinked via cysteine residues, resulting in the formation of a covalently closed ring structure to which the other holocomplex subunits (STAG1 and NIPBL) can bind^13^. Remarkably, our data (Fig.4b,f,g) show that these single-chain cohesin complexes do pass the 30 nm and 200 nm particles during loop extrusion with a similar efficiency (40±6% and 44±16% respectively) to that of wildtype cohesin (Fig. 4e). These data demonstrate that the roadblock passage during loop extrusion does not involve ring opening of the SMC complex.

From systematically investigating the effects of DNA-binding roadblocks on DNA loop extrusion by SMC complexes, we thus conclude that an SMC complex can effectively traverse roadblocks, a finding that even holds for surprisingly large obstacles. Our results directly show that single nucleosomes pose no barrier to condensin-induced loop extrusion, and our finding that RNA polymerases do not block DNA loop extrusion is consistent with Hi-C experiments on bacterial condensin^39^, with the ∼14 s stalling times that we observed similar to the estimated *in vivo* stalling time of ∼10 s for highly transcribed operons.^39^ Our data have important consequences for understanding the activity of SMCs in cells, since the ability of SMC complexes to overcome very large protein complexes on DNA has obvious advantages in its processing of cellular chromatin. Vice versa, the finding that SMC complexes are largely unobstructed by physical barriers along the DNA indicates that SMC barriers such as CTCF must function according to other principles, e.g. be governed by biochemical interactions, as was recently proposed for how CTCF blocks cohesin by binding to the SA2-SCC1 subcomplex of cohesin^21^.

The very large size of nanoparticles that can be accommodated in loop extrusion has also significant consequences for the mechanistic modeling of the SMC motor proteins, as it indicates that SMC complexes can readily bypass obstacles that are clearly larger than its ∼35 nm ring size. Covalent linking of the interfaces of cohesin did not affect its passing ability, showing that a temporary opening of the ring structure is not needed for the loop extrusion process and roadblock traversals. This has direct implications for our understanding of the topology through which the SMC complex interacts with DNA (cf. Fig.4h for a visual explanation of the possible topological modes). Early molecular models for loop extrusion suggested that the SMC ring topologically embraces the DNA upon loading and during DNA loop extrusion.^31,40^ More recently, also a pseudotopological^11,41,13^ and nontopological loading^13,42^ of the DNA were considered. The data from the current study provide unambiguous evidence for a nontopological mode for DNA loop extrusion, where DNA must bind externally to the SMC complex. These finding call for the development of new mechanistic models for the SMC motor functionality that underlies DNA loop extrusion.

## Acknowledgments

We thank E. van der Sluis and A. van den Berg for protein purification; H. Sanchez for plasmid design, N.H. Dekker for supervision of H. Sánchez and T.L., and M. Tisma and A. Katan for discussions.

## Author contributions

B.P., E.K., J.T., C.D. designed the experiments. B.P., R.B. performed most single-molecule experiments and analyzed the data. E.K. performed experiments on dCas9 and RNA polymerase. I.F.D., B.D. purified labeled cohesin and single chain cohesin, under supervision of J.M.P. J.T. prepared the DNA, labeled histone octamers, and reconstituted nucleosomes. T.L. purified histone octamers. W.W. imaged and analyzed functionalized particles in electron microscopy. J.K.R. did AFM imaging.. C.D. supervised the project. All authors contributed to the writing of the manuscript.

## Data and materials availability

All data are available in the manuscript or the supplementary material.

