## Supplementary material for "SMC complexes can traverse physical roadblocks bigger than their ring size": supplimetary info

**This PDF file includes:**

Materials and Methods
Tables S1 to S3
Figs. S1 to S10
Caption for Movies S1 to S6

**Other Supplementary Materials for this manuscript include the following:**

Movies S1 to S6

### Materials and Methods

#### DNA and condensin preparation

Condensin holocomplex from *S. cerevisiae* was purified using our previously published protocol.[1]Lambda DNA containing a biotin on both ends, was made by hybridizing and ligating short oligonucleotides containing 5' phosphate and 3' Biotin on the single-stranded DNA ends of Lambda DNA (NEB, N3011S). For this we used oligonucleotides, JT41 (P-GGGCGGCGACCT-Bio) and JT42 (P-AGGTCGCCGCCC-Bio) (IDT). Taq DNA ligase (NEB, M0208L) was used to ligate the oligonucleotide on Lambda, using 10 times molar access of oligonucleotide to Lambda DNA. The mixture was incubated for 10' at 65°C and then 1hour at 50°C in Taq DNA ligase buffer. The biotin-Lambda DNA was then cleaned up from free oligonucleotides and enzymes using an AKTA pure system (Cytiva), with a homemade gel filtration column containing approximately 46ml of Sephacryl S-1000 SF gel filtration media (Cytiva), run with TE + 150mM NaCl buffer at 0.2ml/min. The fractions containing the Biotin-Lambda DNA were aliquoted and stored at -20°C.

#### Nucleosome reconstitution and AFM imaging

Histone octamers were purified and assembled as established by Sanchez et al. (in preparation). pET-Duet.H3(D82C)-H4 and pCDFduet.H2A(K120C)-H2B were made, using site-directed mutagenesis with pfu-Turbo (Agilent) and Dpn1 (NEB), from pET-Duet.H3-H4 with primers TL-CH-13 (GCGCAGGATTTCAAAACCTGTCTGCGTTTTCAGAGCAGCG), TL-CH-14 (CGCTGCTCTGAAAACGCAGACAGGTTTTGAAATCCTGCGC) and pCDFduet.H2A-H2B with primers TL-CH-5 (CAAAACTTGTTGCCATGCAAGTCTGCCAAGACTGC), TL-CH-6 (GCAGTCTTGGCAGACTTGCATGGCAACAAGTTTTG).

Histone proteins were co-expressed in Escherichia Coli strain BL21-codonplus-DE3-RIL (Agilent) from pET-Duet.H3(D82C)-H4 and pCDFduet.H2A-H2B or pET-Duet.H3-H4 and pCDFduet.H2A(K120C)-H2B.[2] Cells were grown to ~OD 0.4 and expression was induced by adding 0.4 mM isopropyl 1-thio-ß-D-galactopyranoside (Santa Cruz Biotechnology Inc), and shaking at 180 rpm for 16h at 18°C. Cells were subsequently sonicated in a Qsonica Q500 sonicator for 2 min with cycles of 5 s on and 5 s off and an amplitude of 40%, in 0.5M NaCl, 20mM Tris-HCl pH8, 0.1mM EDTA, 1mM DTT, 0.3mM PMSF and protease complete inhibitor (Roche). Supernatant containing the Histone complexes were purified on a 5ml Hi-Trap

Heparin column (Cytiva) and eluted with a gradient of 0.5–2M NaCl, 20mM Tris-HCl pH8, 0.1mM EDTA, 1mM DTT. Fractions were analyzed using SDS-Page and further purified on a Superdex 200 increase (Cytiva) in 2M NaCl, 20mM Tris-HCl pH8, 0.1mM EDTA and 1mM DTT, analyzed on a 12% SDS-Page gel, concentrated and then labeled with C2-Maleimide-Alexa647 (ThermoFisher). For labeling, 1 mg/ml Histone-octamers were dialyzed in 50mM MOPS pH7, 2M NaCl. TCEP (Sigma-aldrich) was added to 1mM and 0.2mM C2-Maleimide-Alexa647 was added. This was incubated for 2hours at 21°C, quenched with 1mM DTT and then purified on a Superdex 200 increase column and checked with a SDS-page gel for intact labeled Histone-octamers.

Nucleosome assembly was done using salt dialysis.[3] For every assembly several different ratios of Alexa647-Histone-octamere/DNA were made. 136nM Alexa647-Histone-octamere, ~0.15nM Biotin-Lambda DNA and 2.5-15nM 10kb DNA (ThermoFisher, SM1751) where combined in 2M NaCl, 10mM Tris-HCL pH8, 1mM EDTA and transferred to a slide-A-lyzer mini dialysis device, 3.5K MWCO (ThermoFisher, 88400). It was then dialyzed at 4°C in 100ml of 2M NaCl, 10mM Tris-HCl pH8, 1mM EDTA, 5mM β–mercaptoethanol. We added 10 mM Tris-HCl pH 8.0, 1 mM EDTA, pH 8, 5 mM β–mercaptoethanol at a rate of 0.5ml/min using a peristaltic pump until the buffer reached ~0.4M NaCl. We then transferred the slide-A-lyzer to a beaker glass with 100ml 10mM Tris-HCl pH 8.0, 1mM EDTA, pH 8.0, 5mM β–mercaptoethanol and incubated for another 2hours. Subsequently we analyzed the sample on an agarose gel and checked on a typhoon if labeled histones where comigrating with the DNA and checked proper nucleosome assembly on the AFM.

After treating a mica with 0.0001 % (wt/vol) poly-L-Ornithine for 1.5 min, we rinsed the mica using 3 ml MiliQ water and dried it using $N_2$ gas. Then, the sample of the nucleosome assembly was deposited onto the mica for 1 min and rinsed by 3 ml MilliQ water. After drying the sample using $N_2$ gas, we imaged it using a Bruker Multimode AFM, with a Nanoscope V controller and Nanoscope version 9.2 software.[4] Using Bruker ScanAsyst-Air-HR cantilevers (stiffness : 0.4 N/m, tip radius : 2 nm). PeakForce Tapping mode was used with an 8-kHz oscillation frequency. To reduce sample distortion induced by tip-sample interaction, we used low a peak force set-point value (< 100 pN). To obtain high resolution images, 2.5 × 2.5 $\mu m^2$ scan areas with 2,500 × 2,500 $pixels^2$ were scanned at 0.5-Hz scanning speed. For imaging condensin and cohesin holocomplexes, we deposited 2 nM protein complexes onto a 0.00001 % (wt/vol) poly-L-Ornithine treated mica for 10 s, and the mica was washed using 3 ml MiliQ water and dried using $N_2$ gas. Then, the sample was imaged with the same AFM parameters described above.

We performed image processing of the AFM images using Gwyddion version 2.53 by removing background subtraction and filtering transient noise.[5] Afterwards, we subtracted (planar and line by line) background polynomials after excluding the masked grains of DNA-protein images and applied plane background subtraction. Finally, we de-convoluted the AFM images using an blind tip estimation and surface reconstruction to minimize the tip convolution effect, the widening of images induced by non-zero AFM tip size. To measure the diameter of nucleosomes (fig. S9d,e), we masked each nucleosome complex bound to DNA on the AFM images and performed mean diameter measurement in the Gwyddion software.

**RNA polymerase labeling and binding to DNA**

Wild-type *E. coli* RNA polymerase core-enzyme (α2ββ'ω) with a SNAP-tag and transcription initiation factor $\sigma_{70}$ were purified and expressed as previously described[6]. Purified RNAP was covalently attached to Alexa647-benzylguanne according to the protocol provided by the provider and further purified using a Sepharose 6 gel-filtration column (Cytiva). The labeling efficiency of the core-enzyme was 35 %. RNAP holo-enzyme was reconstituted by incubating labeled RNAP with $\sigma_{70}$ in 1:10 ratio in a buffer containing 10 mM Tris-HCl, pH 8, 150 mM NaCl, 0.1 mM EDTA, 5 µM $ZnCl_2$, 1 mM DTT, 5 % glycerol. Prior to anchoring λ-DNA onto the surface , 1 nM λ-DNA was incubated with 10 nM RNAP holo-enzyme in imaging buffer (50 mM TrisHCl pH 7.5, 50 mM NaCl, 2.5 mM $MgCl_2$, 1 mM DTT, 5%(w/v) D-dextrose, 2 mM Trolox, 40 µg/mL glucose oxidase, 17 µg/mL catalse) for 5 min at room temperature.

**dCas9 binding to DNA**

gRNA was obtained by annealing a mixture of Alt-R Crispr-Cas9 tracrRNA and crRNA (IDT) at $90^0$ C for 1 min. To prepare dCas9/gRNA construct, 120 nM of dCas9-Snaptag and 1.2 µM gRNA were incubated in NEBuffer3.1 for 30 min at $37^0$ C for 30 min. The dCas9/gRNA construct were stored in aliquots at $-80^0$ C until further use. crRNA target sequences were chosen using the Chopchop webtool (https://chopchop.cbu.uib.no/).

The dCas9/gRNA was attached to the target sequences on λ-DNA by incubating 120 pM biotinylated λ-DNA and 12 nM dCas9/gRNA in NEBuffer3.1 for 45 min at room temperature. We call this construct "λ-DNA/dCas9" in short. To add a label on dCas9 attached to λ-DNA, 100 nM SNAP-Surface Alexafluor-647 was incubated with the "λ-DNA/dCas9" for 45 min at room temperature.

**Table S1.** Target sequences for binding dCas9 to lambda DNA

| ChopChop Rank | Target sequence including PAM | Lambda DNA location | Efficiency by ChopChop |
|---|---|---|---|
| 16 | AAGTGATGCGAAAAAAACAGCGG | seq:10643 | 75.57 |
| 5 | TGTATGAAGATTCACAACCGGGG | seq:19879 | 79.29 |
| 3 | GAAATCCACTGAAAGCACAGCGG | seq:31232 | 81.03 |
| 12 | GCTTGGAACTGAGAAGACAGCGG | seq:38428 | 76.97 |

**Gold and polystyrene nanoparticle functionalization**

For each size, 100 µL of gold nanoparticles (Nanopartz) stock was incubated on a shaker at $4^0$ C overnight with SH-PEG-COOH and SH-PEG-NH2 with concentrations given on the table below. Unbound PEG was removed by dilution (15x) with MilliQ water, centrifugation (speeds in the table), and removal of the liquid supernatant after centrifugation. The dilution and centrifugation step was repeated at least five time to get rid of the unbound PEG. The PEG functionalized gold nanoparticles were incubated with 10 mM Alexa647 NHS and 0.1 mM Benzyl Guanine NHS (BG-NHS) in 50 mM borate buffer at $4^0$C overnight. The unbound dye and BG-NHS was removed by dilution with MilliQ water, centrifugation and removal of the supernatant. The dilution and concentration step was repeated for at least five times to remove most of BG-NHS. We estimate around 1 to 10 BG and around 100 Alexa647 molecules per nanoparticle. The dye functionalized gold nanoparticles were stored at $4^0$C for further use.200 nm amine functionalized polystyrene particles (Nanocs) were incubated with 10 mM SMCC (succinimidyl4-[N-maleimidomethyl]cyclohexane-1-carboxylate) in borate buffer pH 8 for 30 mins in ice bath which leaves maleimide groups on the surface for further reaction with thiol groups. The unreacted SMCC were removed by dialysis. The maleimide functionalized polystyrene particles were immediately functionalized with Alexa647 and BG-NHS as per the Table-2.

Functionalized gold nanoparticles were incubated with 0.5 mg/mL BSA for 5 min and then mixed with λ-DNA/dCas9 for 45 mins at room temperature at a ratio of 10:1 (particles : DNA). λ-DNAs with nanoparticles bound on them were then used to anchor onto the functionalized glass substrate.

**Table S2**. concentration of functionalization groups for particle functionalization

| Size/nm | SH-PEG-COOH/mM | SH-PEG-$NH_2$/mM | Alexa647-NHS/mM | BG-NHS/mM | Centrifugation speed / g |
|---|---|---|---|---|---|
| 14 | 10 | 5 | 10 | 0.1 | 20000 |
| 21 | 10 | 2.5 | 10 | 0.1 | 10000 |
| 29 | 10 | 1 | 10 | 0.1 | 2500 |
| 50 | 10 | 0.4 | 10 | 0.1 | 1100 |
| 125 | 10 | 0.2 | 10 | 0.1 | 400 |
| 200 (polystyrene) | 10 | 1 | 10 | 0.1 | 400 |

**Nanoparticle size measurement from TEM images and FCS**

Hydrodynamic diameters were estimated through FCS measurement. Fluorescence time traces and autocorrelations were recorded in a Picoquant MicroTime 200 confocal microscope. The functionalized gold nanoparticles with Alexa647 on their surface were excited with 640 nm laser with powers giving to 10000 photons per second in average at the detector. The concentration of the particles was kept around 1 nM. The autocorrelation curves (fig. S4) were fitted with

$$G(\tau) = \left[1 - \tau + \tau e^{\frac{-\tau}{\tau_T}}\right] \frac{1}{N(1-\tau)\left(1+\frac{\tau}{\tau_D}\right)} \sqrt{\frac{1}{1+\left(\frac{\tau}{\tau_D}\right)\left(\frac{\omega_0}{\omega_z}\right)^2}} \ ,$$

where τ is the lag time, $\tau_T$ is the triplet blinking time of the Alexa647 fluorophore, N is the number of molecules in the detection volume, $\omega_0$ and $\omega_z$ are the lateral and longitudinal width of the point spread function of the microscope, and $\tau_D$ is the diffusion time of the particle.

$\omega_0$ and $\omega_z$ were determined by calibrating the microscope with the known diffusion coefficient (3.3 $\mu m^2 s^{-1}$) of Alexa647. The hydrodynamic diameter ($2R_h$) was calculated from that as $R_h = \frac{k_B T}{6\pi\rho D}$, where $D = \frac{\omega_0^2}{4\tau_D}$ and T is the temperature, $\rho$ is the viscosity and D is the diffusion coefficient.

Functionalized gold nanoparticles were incubated with 0.5 mg/mL BSA for 5 min and then mixed with λ-DNA/dCas9 for 45 mins at room temperature at a ratio of 10:1 (particles : DNA). These λ-DNAs with nanoparticles bound on them were then used to anchor onto the functionalized glass substrate.

We performed TEM and SEM imaging to verify the sizes of the nanoparticles. The functionalized nanoparticles were diluted 1:20 times with MilliQ and then deposited on carbon grids coated copper TEM grids (Ted Pella) using a JEOL JEM-1400 transmission electron microscope with an acceleration voltage of 120 keV. For particles larger than 100nm, the nanoparticles were deposited on a copper tape and imaged using a FEI Helios G4 CX in SEM mode at 10KeV.

**Single molecule loop extrusion assay**

Flow cells were prepared with PEG/PEG-biotin passivated glass slides as described previously[1]. The channels in the flow cell were incubated with 100 nM streptavidin in Tris20 buffer (40 mM TrisHCl pH7.5, 20 mM NaCl, 0.2 mM EDTA) for 1 min. The unbound streptavidin was washed with Tris20 buffer. The surface was further passivated by incubating the channel with 0.25 mg/mL Bovine Serum Albumin (BSA) in Tris20 for 10 min. 30 µL of 5 pM of λ-DNA with the roadblock on the DNA and 20 nM SxO in Tris20 was flowed into the channel at 4 µL/min. The free DNA in the channel were removed by flowing 100 µL Tris20. The buffer in the channel was replaced with imaging buffer containing 100 nM SxO. The imaging buffer scavenges oxygen and lengthens the observation time of the fluorophores by reducing photo bleaching.

We visualized the DNA and roadblocks in a home built *H*ighly *I*nclined and *L*aminated *O*ptical sheet (HILO) microscope[7] with a 60x oil immersion, 1.49NA CFI APO TIRF (Nikon) objective. HILO allows excitation of molecules in a thin sheet of light that is 1-10 µm above the glass substrate; hence reducing background from free floating dyes in the solution and increasing signal-to-noise ratio of the SxO on DNA and Alexa647 labeled roadblocks. The SxO on DNA was excited with 0.1 W/cm$^2$ 561 nm laser light and single Alexa647 on proteins were excited with 100 W/cm$^2$ 637 nm in alternative excitation (ALEX) mode to simultaneously visualize the DNA and roadblock proteins. The gold nanoparticles with 10 to 100 Alexa647 molecules on the surface were excited with 10 to 100 W/cm$^2$ 637 nm laser light. The two lasers were illuminated alternatively with exposure time of 100 ms allowing simultaneous observation of the DNA and the roadblocks.

**Data analysis**

Fluorescence images were analyzed in a custom written software in python programming language.[8] The images were denoised using a machine-learning-based method called “Noise2void”, as published before[9].

Fluorescence intensities along the DNA axis were averaged with 5 pixels on both side of the axis to obtain a kymograph (e.g. Fig. 1d). Kymographs for the roadblock were obtained with the same axis as the DNA. The intensities of the kymographs was corrected such that non-DNA part of the kymograph had zero intensity. Each vertical line on the kymograph represent a snapshot of the DNA at one time point. Peaks on each line were obtained using the "find_peaks" algorithm in scipy[10] which finds all the local maxima by comparing the neighboring values. The intense peaks from all the peaks obtained were selected with >50% threshold of the peak prominence (relative peak intensities). The peak position represents the loop positon and the loop intensity ($Int_{loop}$) was determined by summing intensities from 7 pixels around the peak position. The intensities of regions ahead ($Int_{ahead}$) and behind the loop($Int_{behind}$), and loop-to-roadblock ($Int_{loop\ to\ roadblock}$) were determined. The intensities were converted to DNA bases as follows:

- DNA size in the loop (bp), $I_{loop} = \frac{Int_{loop} \times 48502}{Total\ DNA\ intensity}$
- DNA size ahead of the loop (bp), $I_{up} = \frac{Int_{ahead} \times 48502}{Total\ DNA\ intensity}$
- DNA size below the loop (bp), $I_{down} = \frac{Int_{behind} \times 48502}{Total\ DNA\ intensity}$
- DNA size roadblock to and including loop (bp), $I_{dist} = \frac{(Int_{loop} + Int_{loop\ to\ roadblock}) \times 48502}{Total\ DNA\ intensity}$

Mean square displacement of the roadblock was obtained from the positions of the roadblock at different times as below:

$$MSD(t) = \frac{1}{50} \sum_{t}^{t+50} |x(t+1) - x(t)|^2 ,$$

where the number 50 denotes the window size of 50 frames. The loop kinetics were smoothed using the Savitzky-Golay method with a 2$^{nd}$ order polynomial with a moving window of 50-100 points.

The result of the encounter between a roadblock and a SMC complex was inferred from both the kinetics and MSD traces. The event was considered an encounter when a roadblock moved towards a loop during loop extrusion and colocalized with the loop. The encounter was inferred from the kymograph. We categorized the results of the encounters into three options: i) the roadblock ended up inside the loop, corresponding to "passing", ii) it remained at the stem of the loop corresponding to "blocking", and iii) a transient pause occurred at the stem of the loop followed by passing into the loop, corresponding to "stalling and passing". An increase in both loop size and MSD after the encounter indicated passing. A constant loop size and MSD upon encounter indicated blocking. An increase in loop size and a constant MSD was also observed occasionally (particularly for cohesin) and was counted as blocking. A constant loop size and MSD, occurring for more than a second, followed by an increase in both corresponded to stalling and passing. Dissociation of roadblock upon encounter with SMC complex was observed only very rarely.

##### Human cohesin and single-chain cohesin experiments

Recombinant human cohesin$^{STAG1}$, NIPBL-MAU2, STAG1 and single-chain cohesin were expressed and purified as described previously.[11] Recombinant cohesin was diluted to 10 nM and NIPBL-MAU2 was diluted to 50 nM in ice-cold storage buffer (25 mM sodium phosphate pH 7.5, 150 mM NaCl, 50 mM imidazole, 5 % glycerol) on ice and briefly mixed by pipetting. Cohesin and NIPBL were then immediately added to imaging buffer (40 mM Tris pH 7.5, 50 mM NaCl, 2.5 mM $MgCl_2$, 5% glucose, 0.25 mg/ml BSA, 1 mM DTT, 0.05% Tween20, 2 mM ATP) supplemented with the oxygen scavenging buffer as described above, equilibrated to 37$^0$C. Imaging buffer supplemented with protein was then immediately introduced in the flow cell at 10 – 30 pM final concentration with a 4-fold molar excess of NIPBL-MAU2 at high flow rate (20-40 µl/min) for 30 sec and flow was turned off to observe loop extrusion events.

Single-chain cohesin was crosslinked by incubation with 0.23 mM BMOE for 10 min on ice. The reaction was quenched by the addition of DTT to 10 mM. For crosslinking controls, BMOE was either omitted or BMOE was quenched with DTT before addition of single-chain trimer (fig. S5a). Crosslinked single-chain cohesin was then incubated with 12-fold molar excess of recombinant STAG1 on ice for at least 30 min (up to 8h). Single-chain cohesin was then added to imaging buffer at 10-30 pM with 4-fold molar excess of NIPBL-MAU2 as for recombinant human cohesin. To check the crosslinking efficiency, ~ 500 fmol of non-crosslinked and crosslinked cohesin were run on a 6% Tris-Glycin protein gel and stained by Coomassie. All experiments with recombinant human cohesin and single-chain cohesin were performed at 37$^0$C.

##### Analysis of cohesin roadblock encounters

Loop extrusion by recombinant human cohesin and single-chain cohesin has been reported to be two-sided,[11,12] in contrast to loop extrusion by yeast condensin that extrudes DNA asymmetrically (3). This fact makes analysis of roadblock encounters with cohesin slightly more involved compared to encounters of roadblocks with condensin. For example, we found that cohesin has a tendency to start extruding loops close to the roadblock on the DNA (~ 60% of loops initiate at the position on the DNA that colocalizes with a roadblock). For loops which initiate sufficiently far from the roadblock, moving of the loop toward the roadblock makes it unequivocally possible to determine which side of the loop encounters the roadblock. However, this is not possible for loops initiating too close to the roadblock. Furthermore, two-sided extruders may keep extruding DNA into the loop from the non-blocked side, thus making loop growth not a sufficient proxy to classify encounters as passing or blocking.

To carefully dissect roadblock encounters with two-sided loop extruders, we distinguish four scenarios, which are illustrated in Fig. S7b. Blocking of the loop extrusion process (case 1) is demarcated by a colocalized DNA loop and roadblock puncta in the kymograph, by halting loop growth and by a decrease of roadblock MSD during initial loop extrusion, and a continuously low MSD as the roadblock is located at the loop base. This case is analogous to the blocking scenario for one-sided extruders. Alternatively, cohesin loop extrusion may be blocked on the side of the roadblock, but continue to extrude on the other side (case 2), in which case DNA loop and roadblock appear to translocate together in the kymograph. The loop keeps growing, yet the MSD of the particle decreases steadily as the loop grows, which allows assignment of the event as blocking, rather than passing as would be judged from the growing loop intensity on its own. A similar argumentation holds for passing events. In the case where the loop continues to grow on the side of the roadblock (case 3), loop and roadblock translocate together, the loop grows and the MSD shows a characteristic minimum at the moment of encounter. For situations in which the loop grows or slips from both sides (case 4), loop and MSD translocate together in the kymograph, yet not necessarily unidirectionally, and the loop size can both grow and shrink. In these cases, a passing event is characterized by a minimum in the MSD as the loop size either remains constant (loop grows on the roadblock side and shrinks on the other side, effectively keeping the loop size constant; we refer to this as ‘loop diffusion’, see Fig. S6f) or increases. A minimum in MSD might also be associated with a decrease in loop size (slipping of DNA out the loop), which releases tension in the DNA, in which case the event cannot be attributed as ‘passing’. Thus the combination of a minimum in MSD and a kymograph with a constant or rising loop size is used as a proxy to distinguish passing and blocking events.

**Table S3**. Key Resources

| REAGENT or RESOURCE | SOURCE | IDENTIFIER |
|---|---|---|
| **Lambda DNA** | New England Biolabs | Cat# N3011S |
| **tracrRNA, crRNA** | Integrated DNA Technologies | NA |
| **dCas9 (SNAP-tag)** | New England Biolabs | Cat# M0652T |
| **1×NEBuffer3.1** | New England Biolabs | Cat# B7203S |
| **Alexa647 NHS** | ThermoFisher Scientific | Cat# A20006 |
| **Benzyl guanine NHS** | New England Biolabs | Cat# S9151S |
| **Gold nanoparticles** | NanoPartz Inc. | NA |
| **Sytox Orange** | ThermoFisher Scientific | Cat# S11368 |
| **Pfu-turbo** | Agilant | 600410 |
| **DpnI** | New England Biolabs | NEB R0176S |
| **TCEP** | Sigma-Aldrich | 646547-10x1ML |
| **BL21-CodonPlus (DE3)-RIL** | Agilent | 230245 |
| **IPTG** | Santa Cruz Biotechnology | Sc-202185 |
| **Protease complete inhibitor** | Roche | 11836145001 |
| **C2-Maleimide Alexa647** | ThermoFisher Scientific | Cat# A20347 |
| **NoLimit 10kb DNA** | ThermoFisher Scientific | SM1751 |
| **Slide-A-Lyzer mini 3.5K** | ThermoFisher Scientific | 88400 |

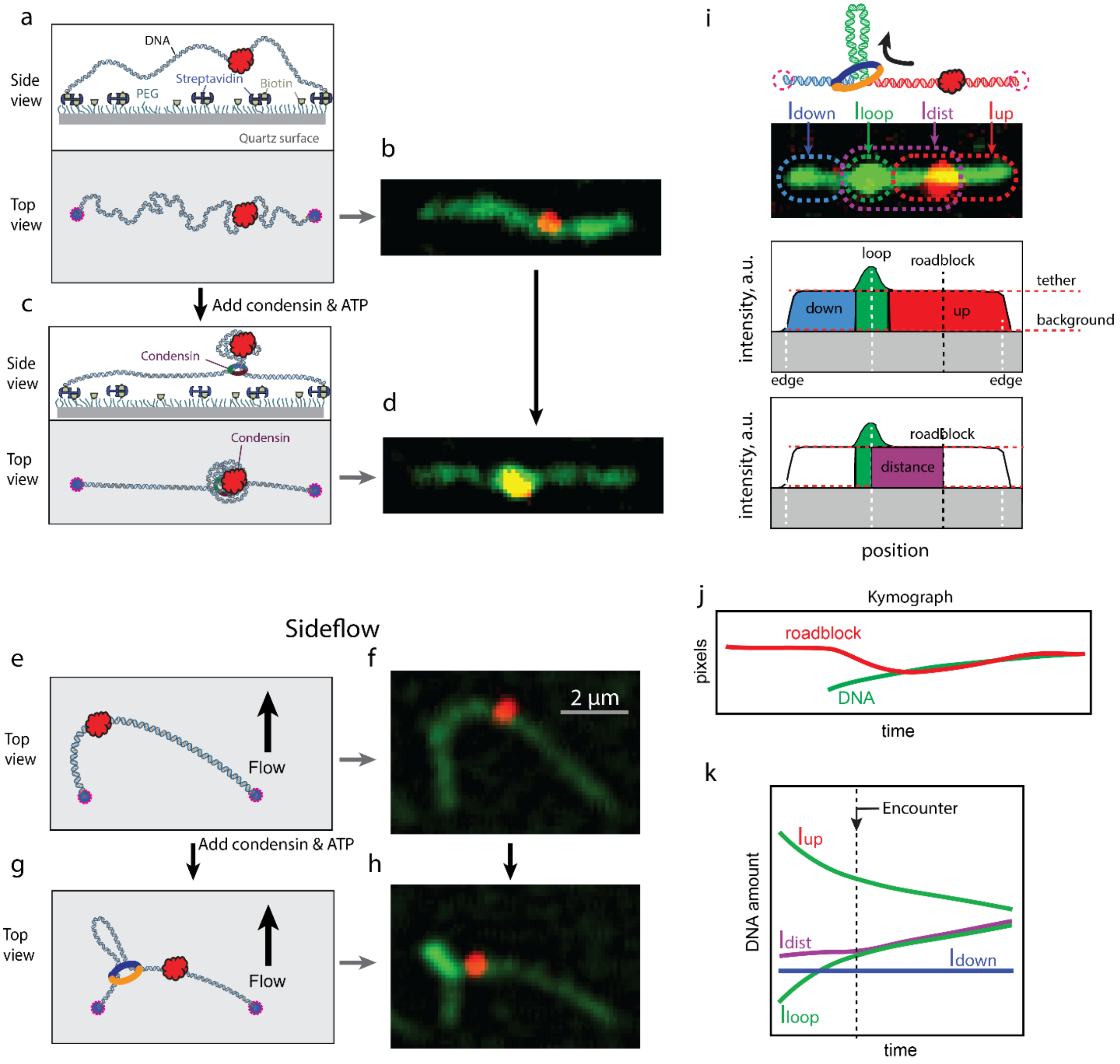


**Fig. S1**. **Experimental Assay**. a) Side view and top view of lambda DNA double tethered on a streptavidin/PEG-coated surface. A roadblock in red is strongly bound to the DNA through dCas9. b) Fluorescence snapshot of the top view of the DNA. c) Side view and top view of an SMC-extruded DNA loop after addition of condensin and ATP. e, f) Top view of the DNA loop extrusion with inplane buffer flow. g, h) Addition of ATP and condensin resulted in a loop and the roadblock moving towards the stem of the loop. i) Representation of intensities of different parts of a loop-extruding DNA molecule with a roadblock. j) Kymograph of the loop position and roadblock position along the axis of the DNA, with x-axis as time and y-axis as pixels. k) Kinetics of intensities of different parts of the DNA as represented in (i).

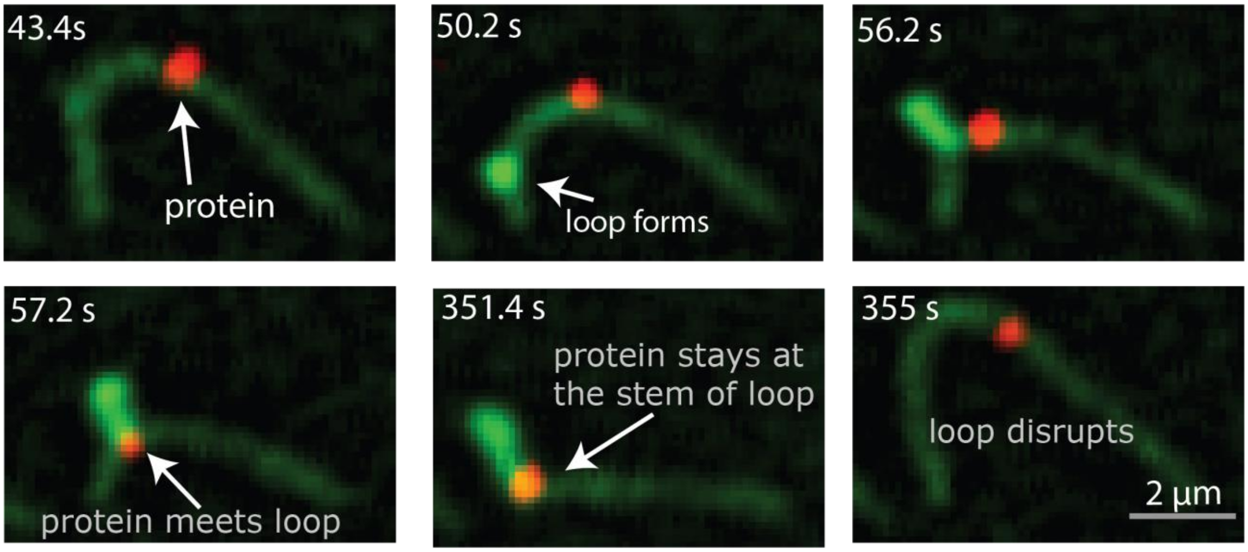


**Fig. S2**. Example of blocking event visualized with a sideflow. The panels show snapshots of condensin extruding DNA with a roadblock protein. The roadblock stayed at the stem of the loop for hundreds of seconds and never passed into the loop, until the loop finally disrupted after 6 minutes.

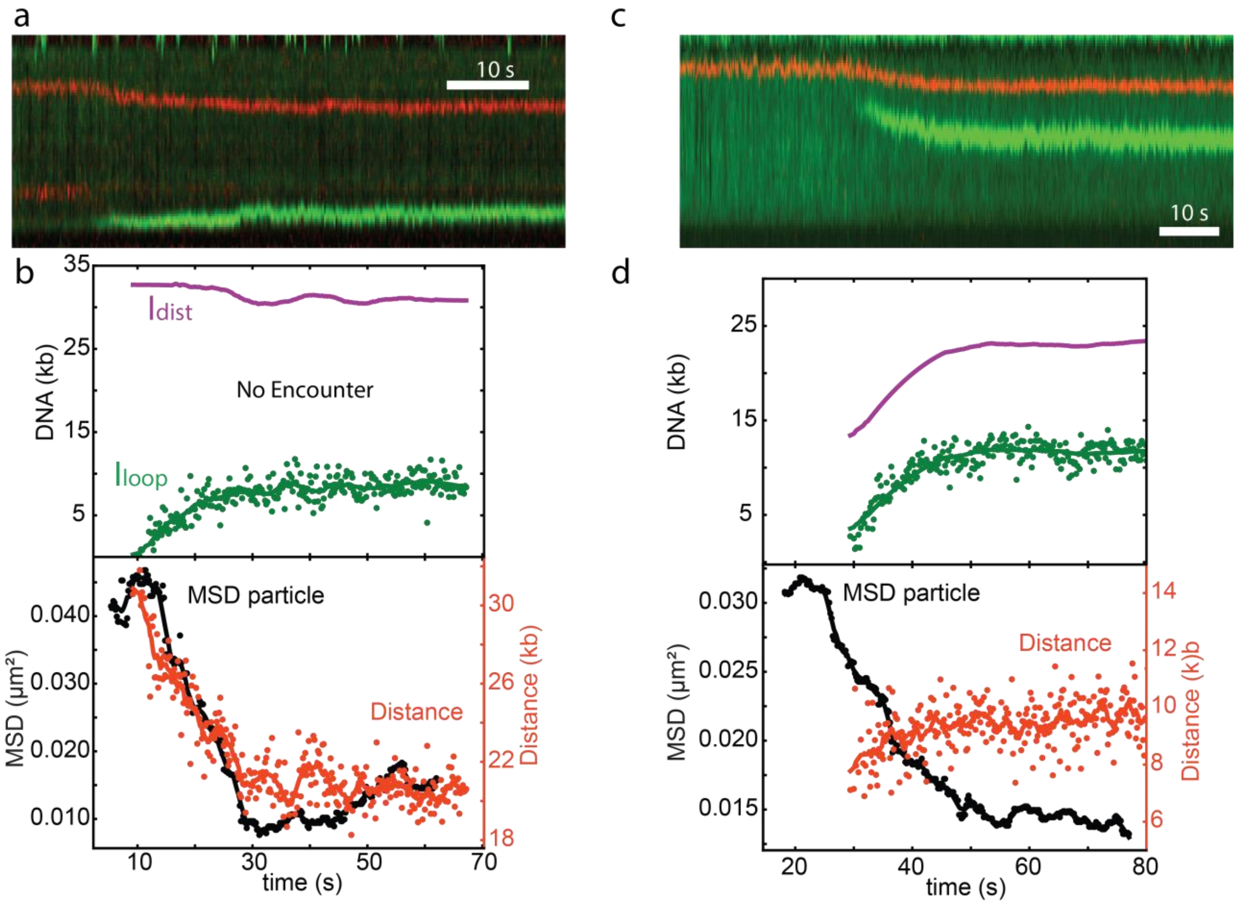


**Fig. S3.** Examples of condensin extruding DNA that has a dCas9 roadblock bound, but where the SMC did not meet the roadblock, and hence the event is not counted as an encounter. a, b) Condensin extruded DNA from the side that contained the roadblock but never reached it due the stalling force that emerged within the DNA. c, d) Example of a roadblock that was on the anchor side of condensin and hence condensin never encountered it.

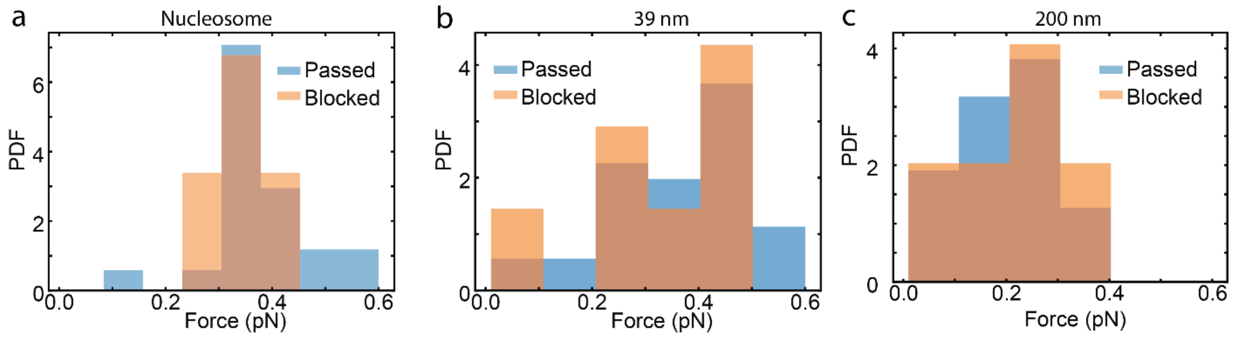


**Fig. S4.** Force histograms for DNA loop extrusion with nucleosomes (a), 39 nm particles (b), and 200 nm particles (c) as roadblocks. Data in blue denote the stalling force for loop extrusion, and data in orange denote the force at which loop extrusion got blocked by the roadblocks. The very similar force values with and without roadblock suggest that blocking is mostly due to lack of slack on the DNA.

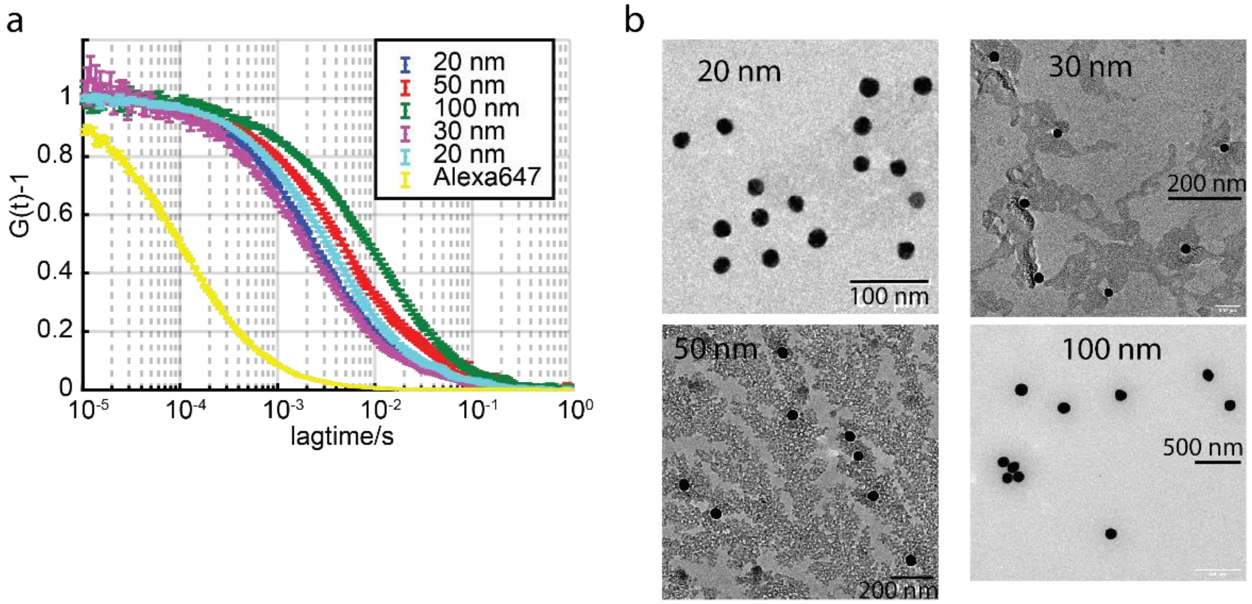


**Fig. S5.** (a) Fluorescence correlation curves for functionalized gold nanoparticles that exhibited a slower decay as the size of the nanoparticle increased. The hydrodynamic diameters that were obtained from these curves (see Methods for details) are denoted in the main text. (b) TEM images of functionalized gold nanoparticles of different sizes showed uniform distributions and absence of aggregates.

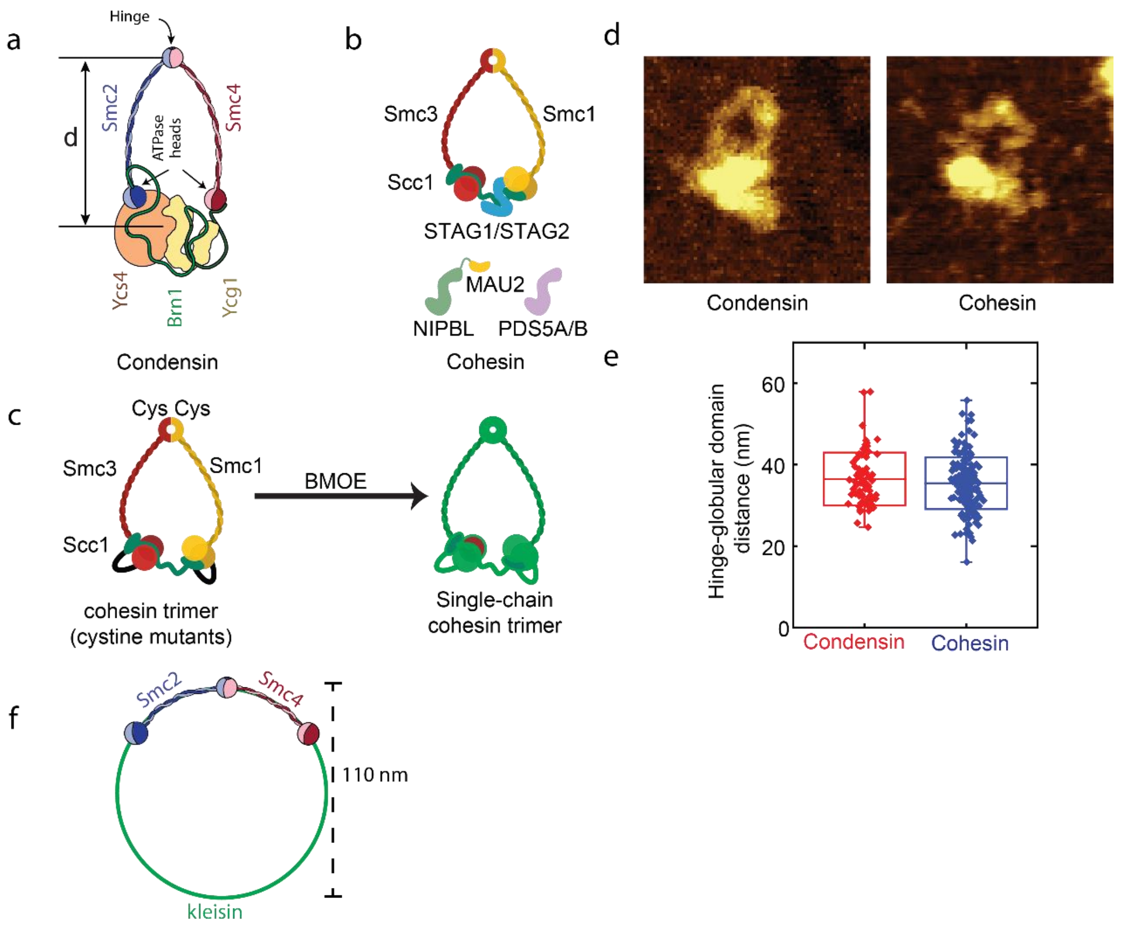


**Fig. S6.** Cartoons of SMC complexes. a) Yeast condensin with its subunits. b) Human cohesin with its subunits. c) Human cohesin trimer fusion protein with cystines at the hinge on both smc subunits that are cross-linked with treatment of BMOE. d) AFM images of O-shaped condensin and cohesin with diameters distributed around a mean value of about 36 nm, as denoted in panel e. The AFM images are 100 x 100 nm. f) Sketch indicating the theoretically maximal circumference that an SMC trimer would be able to attain if the kleisin would be entirely stretched to its full length. The smc2 and smc4 arms are known to be about 50 nm in length each. The kleisin is an unstructured 700 amino acid chain that can, using 0.35 nm/amino acid, be estimated to have a 245 nm theoretical maximum length. This would suggest a maximum possible diameter of about 110 nm for the SMC ring including kleisin in this nonphysiological configuration.

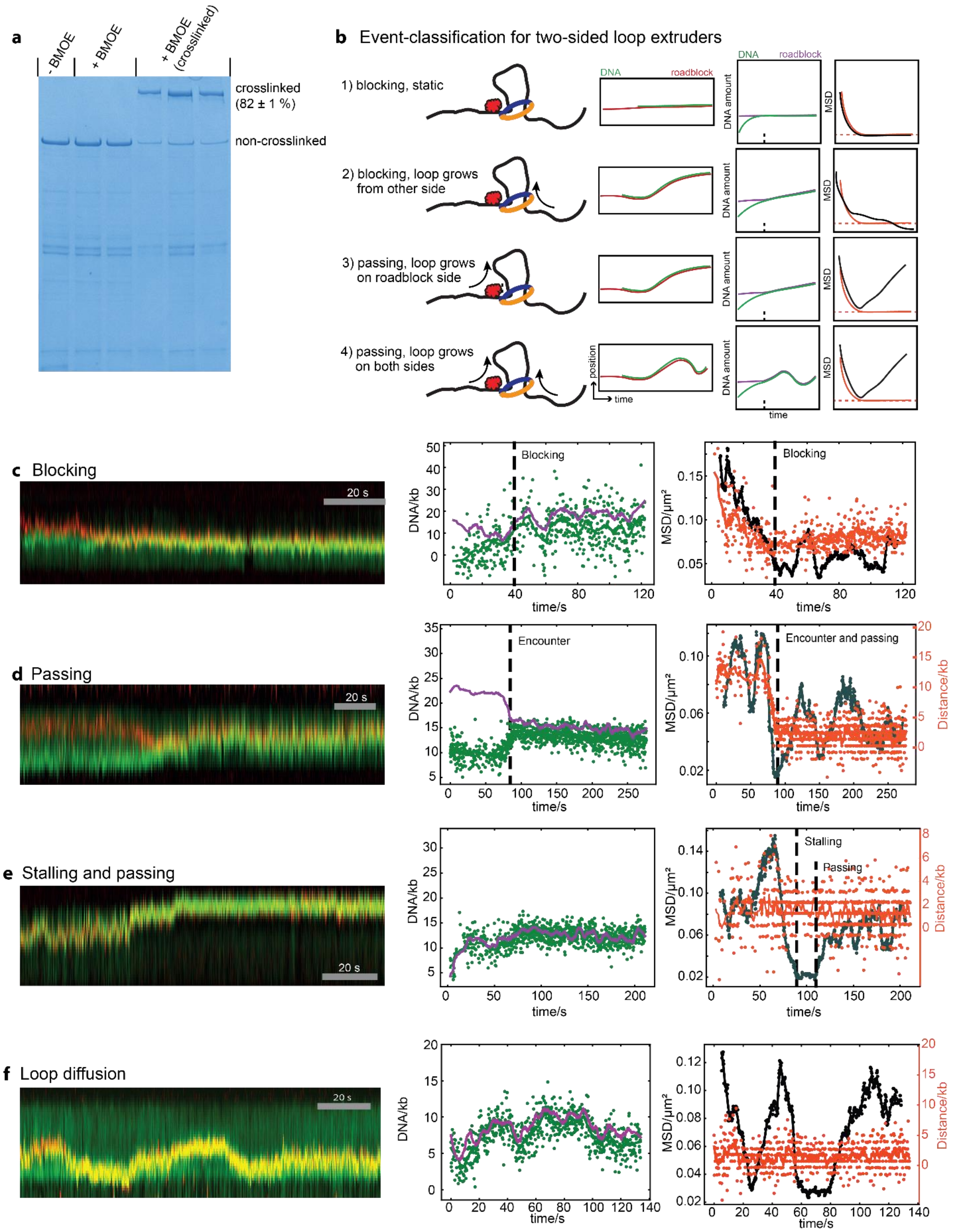


**Fig. S7.** Loop extrusion data for human cohesin on DNA with 30nm gold particles that act as roadblocks. a) Gel shifts of crosslinked cohesin and wildtype cohesin showing a 82% crosslinking efficiency. b)

Illustrative classification of encounter events for two-sided loop extrudes, in particular for cases in which the loop initiates close to or at the roadblock: blocking without further loop growth (1), blocking but loop grows further from the non-blocked side of the DNA (2), passing while the loop only grows from the side of the roadblock (3), and passing while the loop grows and/or shrinks from both sides (4). See supplementary text for an explanation to distinguish these four scenarios. c, d, e) Examples of kymographs and position/MSD traces for loop-extruding cohesin that is getting blocked, passed, and stalled and passed when encountering a roadblock particle, as well as an example where the loop merely diffuses.

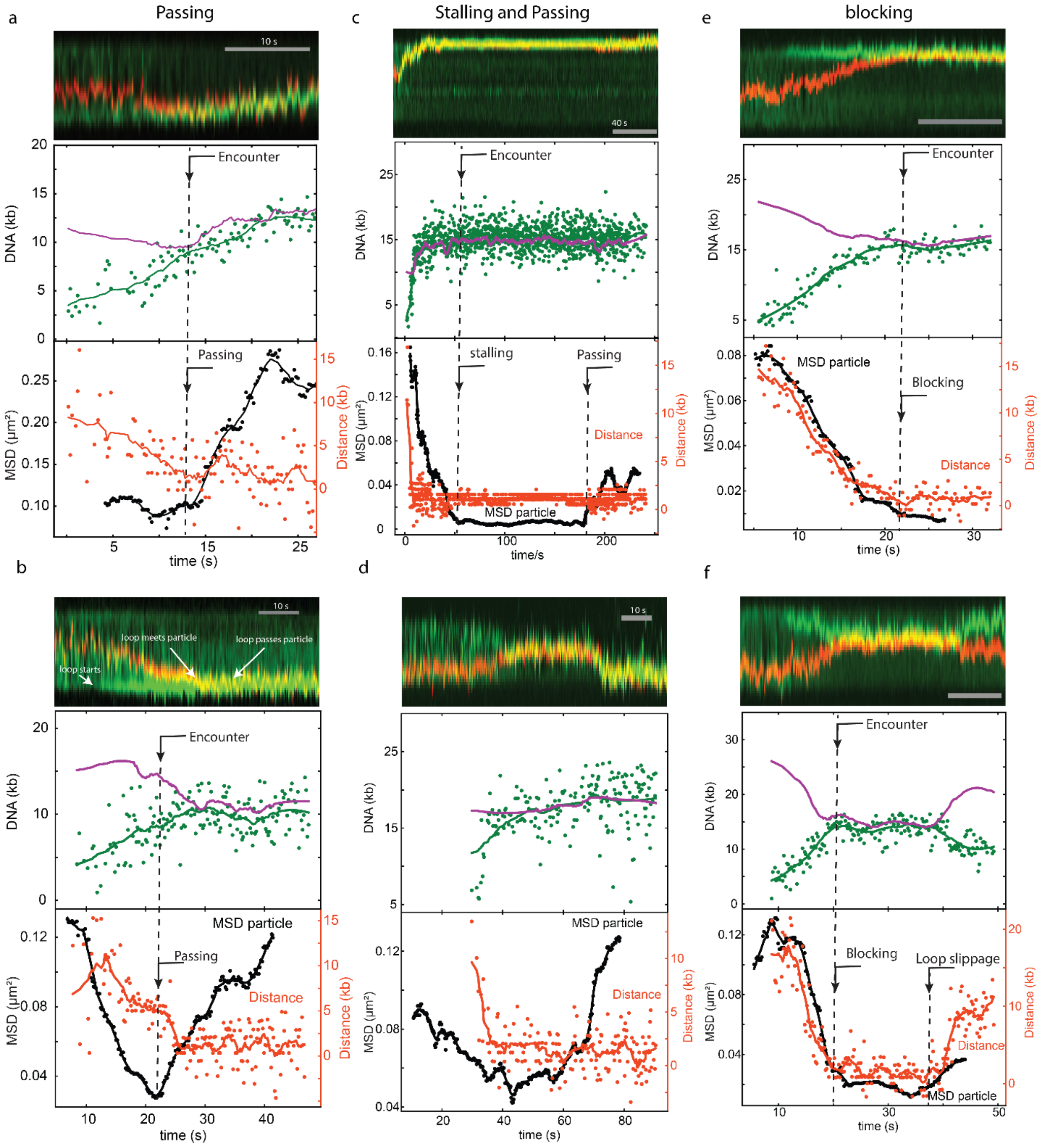


**Fig. S8.** More examples of 200 nm polystyrene beads that act as roadblock for yeast condensin. a,b) Passing event; c,d) stalling and passing; e,f) blocking.

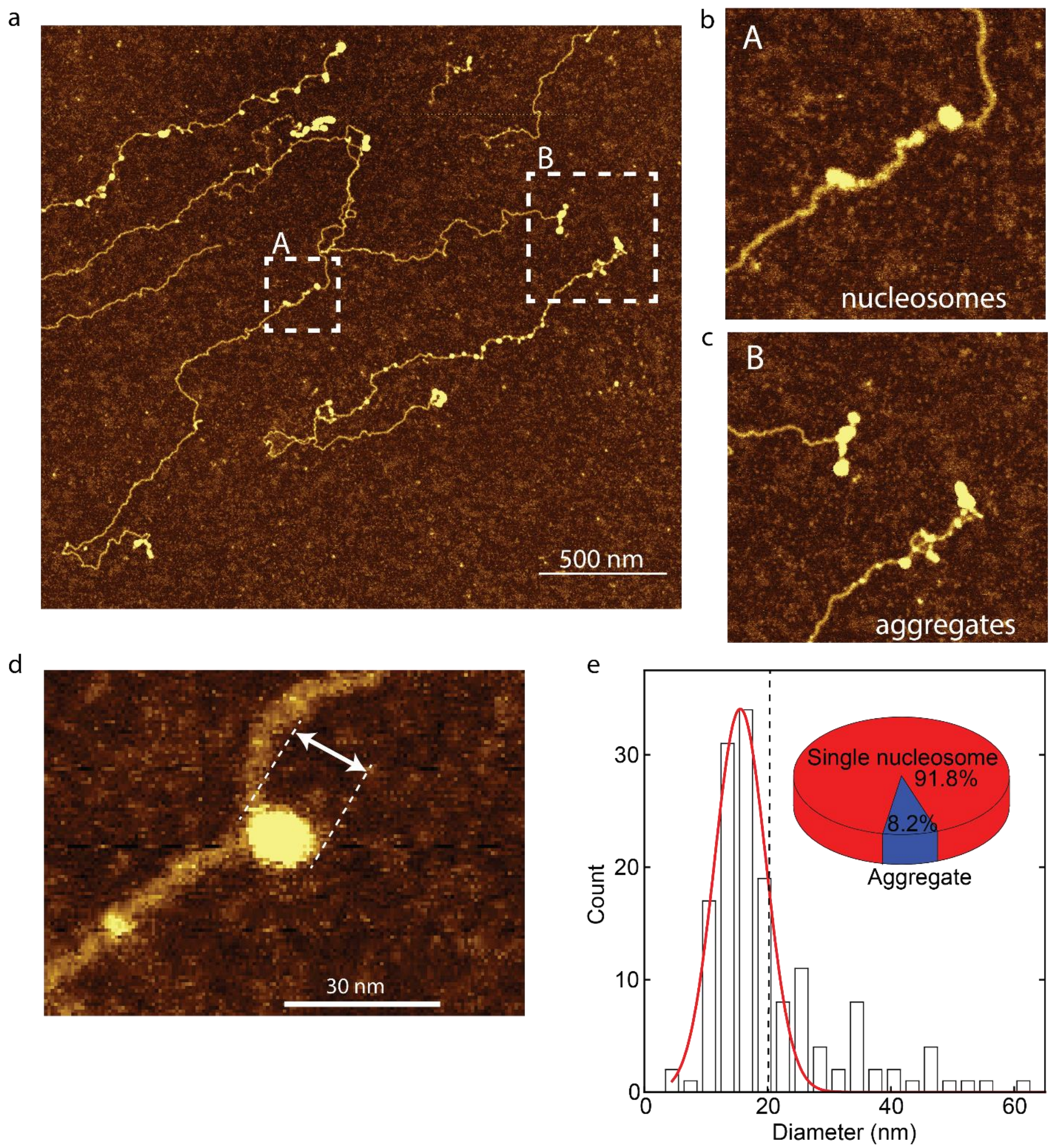


**Fig. S9.** a) AFM image of nucleosomes on DNA. b, c) Zooms of the rectangles shown in (a) for single nucleosomes (b) and some small aggregates (c). (d) Zoom in of a nucleosome with annotation to show the measured diameter. (e) Zoom in of a nucleosome with annotation to show the measured diameter.(e) Histogram of measured diameter of nucleosomes. The solid red line is a single-Gaussian fit yielding a maximum at 15.6 nm. If conservatively, we assign all structures beyond the dashed line as aggregates, we observe that 92% of the nucleosomes are single nucleosomes, see inset, indicating that aggregation was minimal.

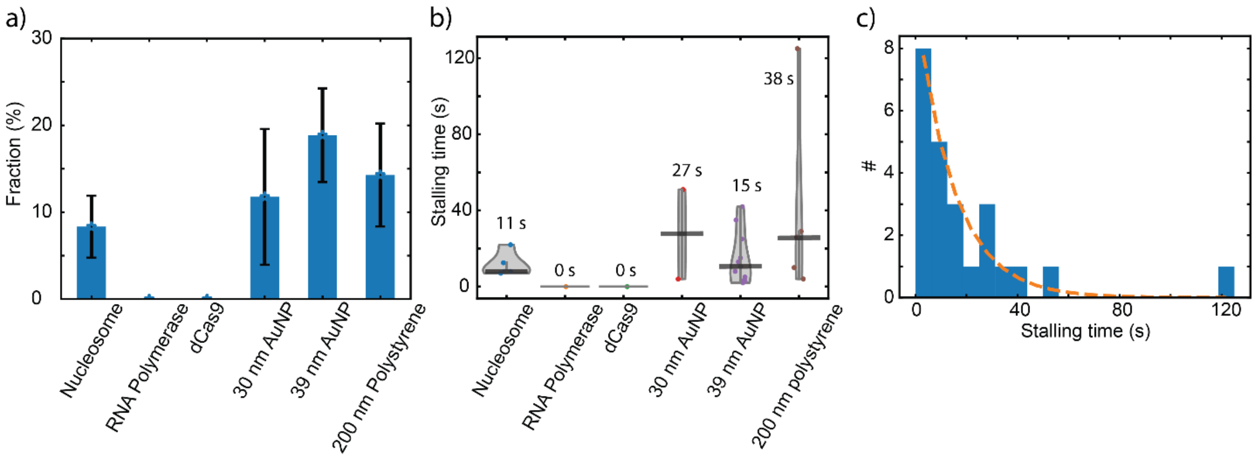


**Fig. S10. Stalling times** a) Fraction of total encounters between condensin and roadblocks that shows stalling and passing. b) Stalling times for different roadblocks. c) Histogram of all stalling times for all the roadblocks shown in (b). Dashed line is an exponential fit with a time constant of 14 s.

**Movie captions**

**Movie S1.** Side flow visualization of condensin passing a dCas9 roadblock during loop extrusion (corresponds to Fig. 1c).

**Movie S2 .** Side flow visualization of condensin blocked by a dCas9 roadblock during loop extrusion (corresponds to fig. S2).

**Movie S3.** Side flow visualization of loop extrusion by condensin with a 39 nm particle as a roadblock (corresponds to Fig. 3f).

**Movie S4.** Loop extrusion by condensin with a 39 nm particle as a roadblock without side flow (corresponds to Fig. 3g). Note that the DNA loop is disrupted at the end of the movie.

**Movie S5.** Loop extrusion by condensin with a 200 nm particle as a roadblock with side flow at the end of loop extrusion (corresponds to Fig. 4c). Note that the DNA loop is disrupted at the end of the movie.
